# Potato juice, a starch industry waste, as a cost-effective medium for the biosynthesis of bacterial cellulose

**DOI:** 10.1101/2021.07.15.452442

**Authors:** Daria Ciecholewska-Juśko, Michał Broda, Anna Żywicka, Daniel Styburski, Peter Sobolewski, Krzysztof Gorący, Paweł Migdał, Adam Junka, Karol Fijałkowski

**Author notes:** **Corresponding author**, E-mail address (K. Fijałkowski).

## Abstract

The unique properties of bacterial cellulose (BC) make it of great interest for numerous branches of industry. Nevertheless, the high cost of the dedicated, microbiological medium used for BC production significantly hinders possibility of widespread use. Searching for an alternative, we turned our attention to potato tuber juice (PJ), a major waste product of the potato starch industry. We verified the possibility of using PJ as a cost-effective, ecological-friendly medium that yielded BC with properties equivalent to those from conventional commercial Hestrin-Schramm medium. The BC yield from PJ medium (>4 g/L) was comparable, despite the lack of any pre-treatment. Likewise, the macro- and microstructure, physicochemical parameters, and chemical composition showed no significant differences between PJ and control BC. Importantly, BC obtained from PJ was not cytotoxic against fibroblast cell line L929 in vitro and did not contain any hard-to-remove impurities. These are very important aspects from an application standpoint, particularly in biomedicine. Therefore, we conclude that using PJ for BC biosynthesis is a path towards significant valorization of an environmentally problematic waste product of the starch industry and can help ultimately lower BC production costs.

**Highlights:**

- Potato juice (PJ) was used as a culture medium for cellulose-synthesizing bacteria.
- PJ was suitable as source of nutrients and did not required any pre-treatment.
- Yield of BC from PJ was equivalent to that obtained from conventional HS.
- PJ-BC did not differ from conventionally produced HS-BC in terms of its properties.
- PJ-BC can be used in the same applications as commercially produced BC.

**Graphical abstract:** 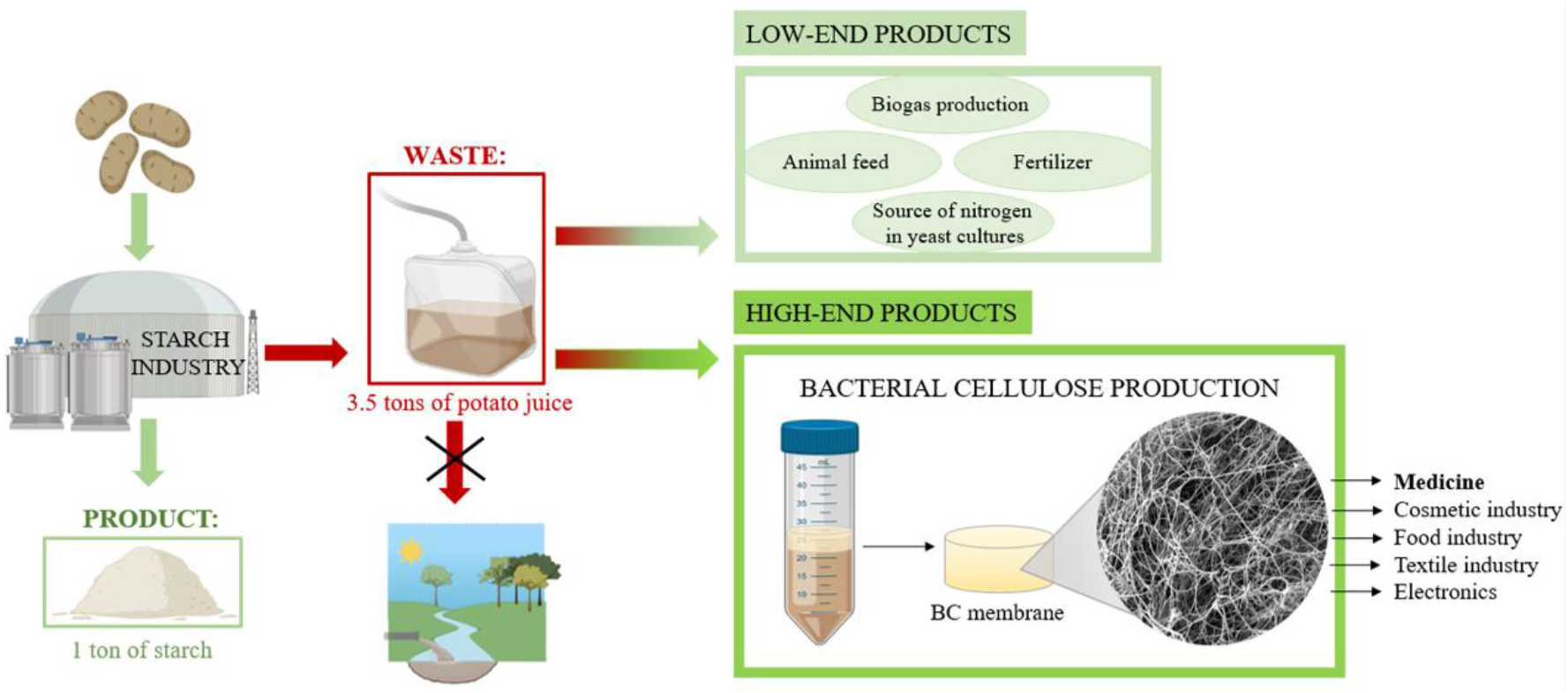

## 1. Introduction

Bacterial cellulose (BC) is a versatile biopolymer, most effectively synthesized by non-pathogenic bacteria, *Komagataeibacter xylinus*. Similar to plant cellulose, BC is a linear polysaccharide consisting of β-1,4-glucan chains (Czaja et al., 2006). However, in contrast to plant-derived cellulose, BC is characterized by flexibility and high chemical purity. Moreover, it does not contain lignins, pectins or hemicelluloses, the presence of which significantly prolongs the purification process of cellulose from plants (Sannino et al., 2009). Additionally, BC has high mechanical strength and water holding capacity thanks to dense fiber structure (Ullah et al., 2019). Importantly, BC can be considered biocompatible and non-toxic, as well as biodegradable, thanks to the activity of cellulase-producing organisms. As a result, BC is safe, not only with regard to industrial applications, but also for environment (Czaja et al., 2006; Sannino et al., 2009). Thanks to the above-mentioned unique properties, BC is versatile, having numerous and diverse applications: in food, cosmetics, pharmaceutical/biomedical, paper, and textile industries (Chawla et al., 2009; Sannino et al., 2009).

In the laboratory setting, BC production is well-established, straight-forward process. Given the proper media and growth conditions, bacteria eagerly produce cellulose membranes. Further, the whole process can be mechanized and, at least partially, automated. However, BC production on an industrial scale requires reduction of costs associated with the relatively expensive medium - typically Hestrin-Schramm (HS) - required for its biosynthesis. From an economic point of view, the less expensive the medium, the higher the potential profits. In this regard, the optimal solution would be the use of an industrial waste, which is dispensable for the manufacturer. This strategy also has additional benefits from an environmental perspective, while re-purposing waste products represents added value. With regard to matter presented in the current study, Kurosumi et al. (2009) examined the possibility of effective BC production (up to 6 g/L of dry BC weight) from various waste fruit juices from oranges, pineapples, apples, grapes and Japanese pears. However, to obtain high yield of the BC, additional nitrogen sources (such as yeast extract) had to be introduced to the aforementioned juices. Otherwise, BC yields were more than 3 times lower (Kurosumi et al., 2009). Lima et al. optimized the method of obtaining BC in the static culture using sisal juice (an agro-industrial residue) as a substrate to produce culture medium supplemented with sugars and yeast extract (Lima et al., 2017). Revin et al. (2018) showed that the use of wheat or whey decoction can enable up to 3 times higher BC yield after 3 days of biosynthesis under dynamic conditions, as compared to conventional media. In turn, Kongruang (2007) proposed a way to reduce the cost of BC production by using coconut and pineapple juices (rich in proteins, carbohydrates and microelements). However, again, these media had to be supplemented with yeast extract (Kongruang, 2007). For the same reasons, Li et al. (2015) investigated the possibility of using wastewater from the process of candied jujube production. The results of their research have shown that such post-production water was inexpensive to obtain; however, it required acid pre-treatment in order to obtain sufficient glucose content. The 3-hour hydrolysis at 80°C resulted in a 58% higher glucose content in the raw material, which translated into high BC yield (Li et al., 2015). Other natural ingredients, reported by several research groups, included hydrolysates of corn stalks or wheat, rice husk pre-treated with enzymatic hydrolysis, fruits and vegetables peels (Abol-Fotouh et al., 2020; Cheng et al., 2017; Goelzer et al., 2009; Hussain et al., 2019; Kongruang, 2007; Kurosumi et al., 2009). Summarizing the above information, it can be concluded, that the main issues with application of natural or waste substrates are related to the mandatory pre-processing stage or insufficient nutritional content. Both of these issues may significantly increase the total cost and extend the BC production process.

With these aspects in mind, we turned our attention to the potato industry. Potatoes are vegetables with high nutritional value and their extensive cultivation in Eastern Europe, China and India translates into high availability and low price throughout the year cycle (Bradshaw and Bonierbale, 2010). At the same time, the potato agro-industry generates substantial waste. Recently, Abdelraof et al. (2019) described their efforts to utilize potato peel waste for obtaining substrates for BC biosynthetic media. To our knowledge, this is only report concerning use of potatoes or potato waste to produce BC. However, the processing of potato peels in order to obtain sufficient sugar content also required use of acidic hydrolysis, with additional reagents and high energy consumption (the hydrolysis process was carried out at a temperature of 100°C). As a result, the potential industrial applicability and cost-effectiveness of this approach is limited.

In our approach, we selected potato juice (PJ), another significant potato agro-industrial waste product. The most prominent producer is the starch industry, which in the European Union produces over 10 million tons of starch and starch-derivatives, and more than 5 million tons of proteins and fibers each year (https://starch.eu/the-european-starch-industry/). During the manufacture of a single ton of potato starch, approximately 3.5 tons of PJ waste is produced. Further, the proper management of PJ and other post-production waste is a significant and unresolved technological problem for this industry (Fang et al., 2011; Grommers and van der Krogt, 2009). Given the growing challenges presented by climate change and other global environmental problems, the proper management of potato agro-industrial waste should be considered a key public policy target for governments and the private sector. Thus far, only the use of potato wastewater to irrigate agricultural fields has been explored; however, it creates problems with soil clogging, loss of water permeability, and water eutrophication (Kot et al., 2020).

Therefore, the goal of the current study was to investigate if PJ can be used as the medium for BC synthesis. We compared the yield, morphology, chemical composition, sorption properties, mechanical strength, and cytotoxicity of BC obtained from PJ-based cultures to those in dedicated HS medium. In this fashion, we demonstrate an environmentally friendly strategy towards valorizing a major agro-industrial waste product.

## 2. Materials and methods

### 2.1. Preparation of culture medium

For the current study, we selected the Tajfun variety potato (average dry matter content: 22%, average starch content: 16%), because it is one of the varieties used by the starch industry. Potato tubers were obtained from Pomeranian-Masurian Potato Breeding Company (Strzekęcino, Poland). Prior to use, the tubers were stored at 4°C, not longer than 4 weeks. Potatoes were washed and peeled and then PJ was obtained using a high-speed juicer (Bosch, MES3500, Germany). Following juicing, the PJ was left at room temperature for 60 min to enable the starch residues settle and then the potato juice was decanted. The decanted PJ was then diluted with distilled water in a 1:1 ratio, sterilized by autoclaving at 121°C, centrifuged for 10 min at 1500 × *g* to remove the remains of precipitated solids, and finally enriched with 1 v/v% of ethanol to yield the final PJ medium **(Fig. 1).**

**Fig. 1.**
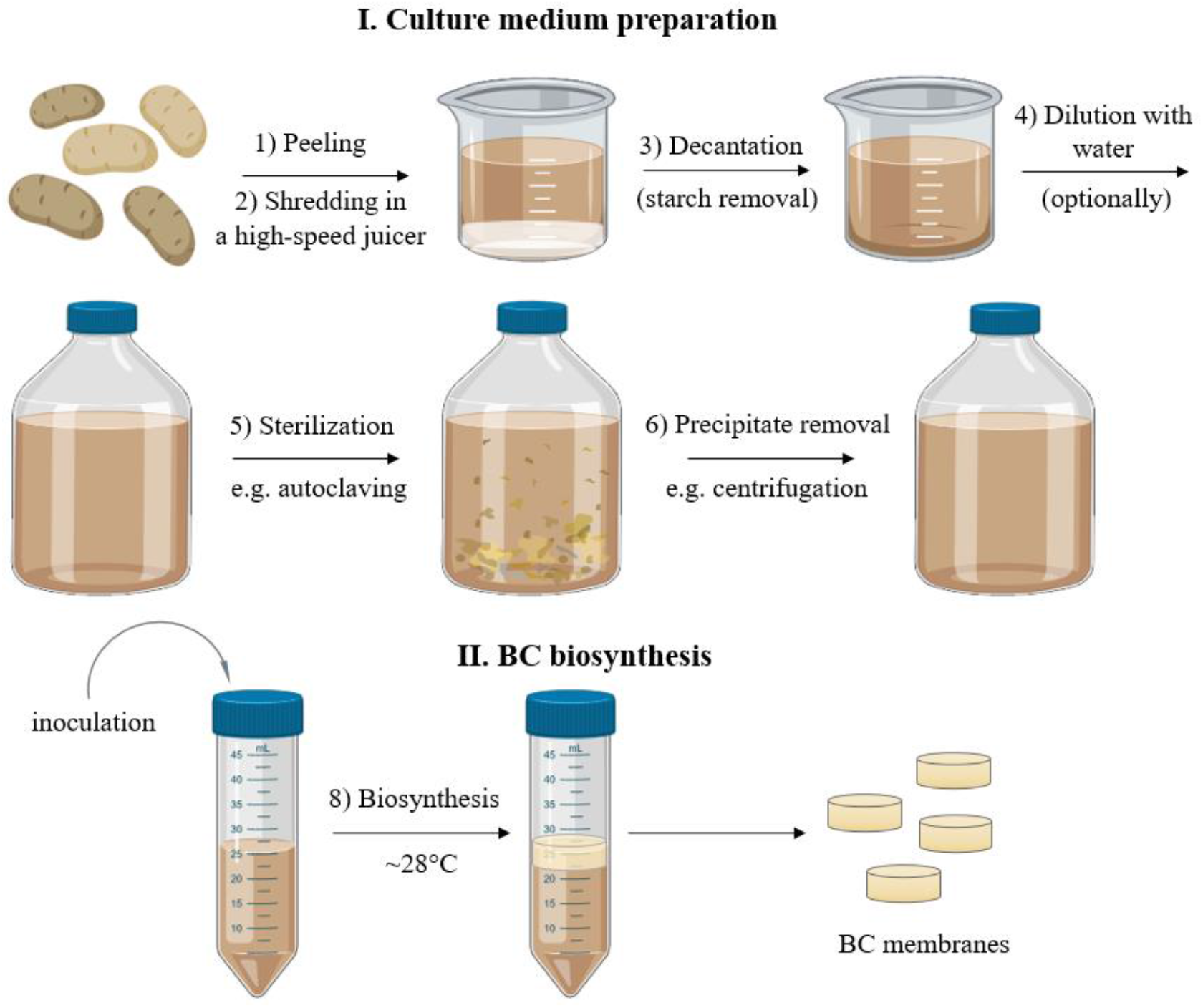
Scheme of the culture medium preparation and BC biosynthesis process. Created using BioRender.com.

### 2.2. Determination of pH, protein, and carbohydrate concentration in PJ medium

The concentration of sugars, including sucrose, glucose, fructose in PJ medium was determined by liquid chromatography - tandem mass spectrometry (LC–MS/MS) technique (1260 Infinity II Series Liquid Chromatograph, Agilent, USA). The InfinityLab Poroshell 120 EC-C18 column (Agilent, USA) with a particle diameter of 2.7 μm equipped with a guard column was used for the chromatographic separation. The mass spectrometer (Ultivo G6465B, Agilent, USA) coupled to the chromatograph was used to detect and to identify the tested analytes. Quantitative analysis was performed based on calibration curves prepared with the use of high purity sugars standards (Millipore Sigma, USA).

The concentration of total protein was measured using the Bradford protein assay with bovine serum albumin as a standard (Bradford, 1976). The starch concentration was measured using the iodine starch method (Sulistyarti et al., 2015). The measurements for both of these assays were performed spectrophotometrically at 595 nm and 615 nm respectively, using an Infinite 200 PRO NanoQuant Microplate Reader (Tecan, Switzerland). The pH of PJ medium was assessed using a laboratory pH meter (Elmetron, Poland).

### 2.3. Microorganisms and culture conditions

For BC production, a reference strain of *Komagataeibacter xylinus* (American Type Culture Collection – ATCC 53524) was used. The bacterial suspension of the cell density equals 2×10^5^ CFU/mL was used to inoculate 25 mL of medium in 50 mL plastic tubes (with 3.8 cm diameter) (Polypropylene Conical Centrifuge Tube, Becton Dickinson and Company, USA). Next, BC synthesis was conducted for 7 days at 28°C. As a control, a standard HS medium (consisting of 2 w/v% glucose, 0.5 w/v% yeast extract, 0.5 w/v% peptone, 0.115 w/v% citric acid, 0.27 w/v% Na2HPO_4_, 0.05 w/v% MgSO_4_·7H2O), enriched with 1 v/v% ethanol, was used.

In order to fully assess the possibility of using the PJ medium and the full spectrum of its potential advantages, several additional experiments were performed in analogous fashion. These included cultures of 7 additional *K. xylinus* strains from American Type Culture Collection (ATCC) - 53582, 23770, 700178, 23768, 23769, 35959, 14851), as well as tests of several HS and PJ media combinations (PJ diluted with distilled water in 1:2 and 2:1 ratio; PJ and HS media without the addition of ethanol; non-centrifuged PJ containing starch solids; HS medium supplemented with the sucrose, glucose, fructose and combination of these sugars at the same concentrations as in PJ). Additionally, we prepared and tested several media containing other ingredients that are agro-industrial wastes, including potato peels, orange peels, beetroot, and apples. The method of preparation of these media was analogous to the one applied for preparation of the PJ medium. Finally, we also tested the use of other culture vessels such as Petri dishes (15×20 mm) - the BC obtained from Petri dishes was also used for evaluation of its mechanical properties.

### 2.4. Determination of BC ield

Starting from the 3^rd^ day of cultivation, BC pellicles synthesized in HS (HS-BC) and in PJ media (PJ-BC) were harvested, purified by treating with 0.1 M NaOH solution at 80°C for 90 min, and then rinsed with distilled water until pH became neutral (6.5 – 7.5). The obtained samples were then weighted on an analytical balance (accuracy 0.0001 g, WTB 2000 Radwag, Poland), dried at 60°C, and weighted again. BC yield was expressed as dry mass (g) of BC/volume of culture medium (L).

### 2.5. Quantification and viability assessment of K. xylinus cells

The quantity and viability of bacterial cells were determined in PJ and HS media (from 1^st^ day of cultivation) and in PJ-BC and HS-BC pellicles (from 3^rd^ day of cultivation) after their enzymatic digestion with cellulase, using alamarBlue Cell Viability assay (ThermoFisher, USA). AlamarBlue Cell Viability Reagent is a ready-to-use resazurin-based solution that functions as a cell viability and metabolic activity indicator. For the digestion, the pellicles were washed in distilled water, transferred to 25 mL of the cellulase solution in citrate buffer (0.05 mol/L, pH 4.8), and incubated with shaking for 24 h at 30°C. Next, the samples consisting of either culture medium or cell suspensions obtained from enzymatic hydrolysis of pellicles were centrifuged for 10 min at 3300 × *g*. The resulting pellets were washed in phosphate buffered saline (PBS, MilliporeSigma, USA), centrifuged again under the same conditions, and restored to their original volume with PBS. The bacterial suspensions (200 μL) were then transferred into wells of 96-well fluorescence microtiter plates (Becton Dickinson and Company, USA) and 20 μL of alamarBlue reagent was added, followed by incubation for 1 h at 30°C, in dark. The fluorescence was measured using microplate fluorescence reader (Synergy HTX, Biotek, USA), using 540 nm excitation and 590 nm emission filters. Sterile PBS was used as the blank. Because *K. xylinus* cells may show different metabolic activity depending on whether they are isolated from the medium or from the BC pellicle, dedicated standard curves were prepared separately for the cells obtained from the culture medium and the BC pellicles. Using these standard curves, the fluorescence data was converted into log CFU/mL.

### 2.6. Assessment of pH, protein, sucrose, glucose, fructose and gluconic acid concentration during BC biosynthesis

Each day of the fermentation process, samples of the culture media were taken and centrifuged for 1 h at 15000 × *g*. The supernatant was then separated from the pellet, filtered (PES 0.22 μm, MiliporeSigma, USA) and if dedicated for the LC-MS/MS analyses, additionally diluted with water (1:1000). All of the measurements were performed as previously described in section 2.2 (*Determination of pH*, *protein*, *and carbohydrate concentration in PJ medium*).

### 2.7. Evaluation of physical and chemical properties of BC obtained from PJ medium

#### 2.7.1. Analysis of microstructure – Scanning Electron Microscopy (SEM)

Purified PJ-BC and HS-BC pellicles obtained after 7 days of culture were fixed in 1% (v/v) aqueous solution of glutaraldehyde for 0.5 h at room temperature and dehydrated using an ethanol dilution series from 10% to 100% (v/v) (5 min at each concentration). Subsequently, the samples were dried at room temperature for 15 min, coated with a 15 nm layer of carbon using a high vacuum carbon coater (ACE 600, Leica, Germany), and imaged with the ZEISS Auriga 60 scanning electron microscope (SEM, Auriga 60, Zeiss, Oberkochen, Germany).

#### 2.7.2. Determination of chemical composition of BC pellicles – Attenuated Total Reflectance Fourier Transform Infrared (ATR-FTIR) spectroscopy

Infrared spectra of purified PJ-BC and HS-BC pellicles obtained from 7 days cultures were evaluated using the ATR-FTIR technique, using ALPHA FT-IR Spectrometer (Burker Co., Germany) with a DTGS detector and the platinum-ATR-sampling module with a robust diamond crystal and variable angle incidence beam. For each BC pellicle 32 scans at 4 cm^-1^ resolution were recorded over the spectral range of 4000 - 400 cm^-1^. Initial spectral data processing was performed using the Spectragryph 1.2 software package.

The crystallinity index was calculated using the ratio of absorbance values for peaks 1430/900 (Cr.R1) and 1370/2900 (Cr.R2). The fraction of the cellulose *Iα* was calculated from ATR-FTIR spectra according to the Eq. 1 (Kataoka and Kondo, 1999). The area of the peaks at *A710* (710 cm^-1^) for *Iβ* and at *A750* (750 cm^-1^) for *Iα* was determined from the spectra deconvoluted using Peakfit software. The following equation was used:

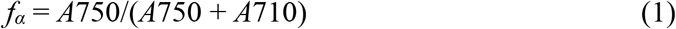

#### 2.7.3. Determination of water swelling ratio

Purified PJ-BC and HS-BC pellicles obtained from 7 days cultures were dried at 60°C for 6 h to remove any water, weighed, immersed in distilled water for 24 h, and weighed again. The swelling ratio as a percent of dry mass (SR%) was then calculated using Eq. 2:

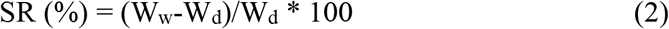

where W_w_ is the weight of the swollen BC and W_d_ is the dry weight of the sample.

#### 2.7.4. Determination of water holding capacity

Purified and dried PJ-BC and HS-BC pellicles obtained from 7 days cultures were weighted, immersed in distilled water for 24 h to obtain maximum absorption level, and weighed again. The ability to hold water was determined using moisture analyzer (Radwag, Poland) at 60°C until the weight of BC was equal to the initial value (dry weight before hydration). Weight measurements were made automatically every 2 minutes. Water holding capacity (WHC) was then calculated using the Eq. 3:

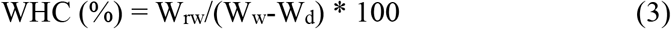

where W_rw_ is the weight of water retained in BC during drying, W_w_ is the initial weight of wet BC, and W_d_ is the dry weight of the sample.

#### 2.7.5. Determination of density

The density of the dry, purified PJ-BC and HS-BC pellicles obtained from 7 days cultures was determined using a hydrostatic balance (XA 52/Y, Radwag, Poland) with methanol as a standard liquid. The weight of samples was measured at room temperature in air as well as in methanol. The sample density was calculated using the Eq. 4 (Ciecholewska-Juśko et al., 2021):

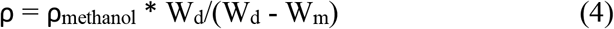

where W_d_ is the weight of the dry sample in the air and W_m_ is the weight of the sample in methanol.

#### 2.7.6. Evaluation of mechanical properties

The tensile tests were performed according to PN-EN ISO 527-1 using Instron Universal Testing Machine (Instron, United Kingdom). For this purpose, purified wet PJ-BC and HS-BC pellicles obtained from 7 days cultures in Petri dishes, that were cut into 2.5 × 10 cm strips. Prior to testing, samples were gently pressed to remove excess water and obtain uniform thickness of 4.5 to 5 mm. Tensile tests were then carried out at speed 10 mm/min, at room temperature. The average values of tensile strength and elongation at break were calculated from the stress-strain curves. All measurements were performed in four repeats.

### 2.8. Cytotoxicity screening

*In vitro* cytotoxicity screening of the purified PJ-BC and HS-BC pellicles was performed using extract and direct contact assays, based on 10993-5:2009 standard using ATCC CCL-1 (L929) murine fibroblasts (passages 10-28), as described previously (Ciecholewska-Juśko et al., 2021). L929 cell line, Dulbecco’s Modified Eagle Medium (DMEM), fetal bovine serum (FBS), L-glutamine, penicillin, streptomycin, and all other cell culture reagents were purchased from MilliporeSigma (MilliporeSigma, USA). All cell culture plasticware and disposables were purchased from VWR (VWR, USA). L929 cells were maintained and cultured in DMEM supplemented with 10% FBS, 2 mM L-glutamine, 100 U/mL penicillin. and 100 μg/mL streptomycin. For all cell culture assays, BC pellicles were sterilized by autoclaving.

#### 2.8.1. Extract assay

For extract assay, 5 pellicles (~10 cm^2^) of each material were placed in a 6-well plate well, covered with 5 mL of growth media, and incubated for 24 h in a cell culture CO2 incubator at 37 °C. For dried HS-BC/PJ-BC films, 5 discs (~10 cm^2^) were covered with 3 mL of media. Finally, just media incubated in the same fashion served as a sham extract, while extracts of medical-grade PCL (CAPA 6430) and nitrile glove served as negative and positive (toxic) controls, respectively. In parallel, a 96-well plate was seeded with 1 x10^4^ L929 cells per well and incubated for 24 h to allow for cell adhesion and spreading. At this point, the media was replaced with 100 μL of each extract, with 6 technical replicates performed per material. The plate was then returned to the incubator and cells were cultured for an additional 24 h, after which cells were examined using inverted light microscope (Delta Optical IB-100). Cell viability was evaluated using resazurin assay. Fluorescence measurements were performed using fluorescent plate reader (Synergy HTX, Biotek, USA) at 540 nm excitation and 590 nm emission. Complete growth media in an empty well was used as a blank. The results were expressed as percent of cell viability relative to control (sham) and were calculated using Eq.5 (Riss et al. 2004):

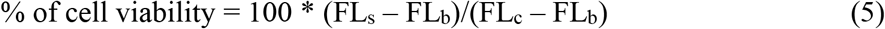

where FL is the fluorescence intensity (arbitrary units) and indexes *s*, *b*, and *c* refer to sample, blank, and control, respectively.

#### 2.8.2. Direct contact assay

For the direct contact assay, 5×10^4^ L929 cells were seeded per well of a 24-well plate and incubated for 24 h to allow for cell adhesion and spreading. After this time, the media was replaced with fresh media and BC pellicles (~2 cm^2^) that had been presoaked in cell culture media were placed directly on top of the cell monolayer (n = 5 pellicles per material). After further 24 h of incubation, pellicles were carefully removed, and cell viability was evaluated as described above using both light microscopy and the resazurin viability assay.

#### 2.8.3. Visualization of fibroblast viability

L929 fibroblasts cultured and treated as for the extract and direct cytotoxicity assays were stained for 15 min. with 3 μL of Syto 9 and Propidium iodide (PI) dyes (ThermoFisher Scientific, USA) 1000-fold diluted in PBS (MilliporeSigma, USA). Next, the dye-containing buffer was removed, and the cells were gently rinsed 3x times with PBS. Images of stained cells were captured using Lumascope 620 (Etaluma, Carlsband, CA, USA) at magnification x20.

### 2.9. Statistical analyses

The data obtained in this study are presented as mean values ± standard error of the mean (SEM). Statistical differences between BC samples obtained in cultures carried out in HS and PJ media were determined by one-way analysis of variance (ANOVA) and Tukey’s post hoc test. The cultures were conducted in triplicate and all experiments were repeated at least three times. Differences were considered significant at a level of p<0.05. The statistical analyses were conducted using Statistica 12.5 (StatSoft, Inc. Tulsa, OK, USA).

## 3. Results and Discussion

### 3.1. Characterization of PJ medium

In the current study, we used potato tuber juice (PJ) as a culture medium for BC biosynthesis. Importantly, the PJ was obtained in the same way as during industrial production of starch. Further, the PJ medium required no pre-treatment process or any additional supplementation with nutrient sources, with the exception of the addition of 1% (v/v) ethanol. Ethanol is typically supplemented into HS media, because it is a stimulating/initiating factor for BC biosynthesis by *K. xylinus* (Chawla et al., 2009).

The lack of necessity of supplementing PJ with additional nutrients arises from the high content of various, easily absorbed nutrients present in potato tubers. While starch is the main component of the dry mass of potatoes (approx. 2%), tubers also contain: 1) approx. 2% of proteins (a source of amino acids for bacterial cells), 2) 0.3 to 0.6% of sugars (a carbon source), and 3) a wide range of vitamins (e.g. A, C, B1, B2, B6) and minerals (e.g. potassium, phosphorus, magnesium, calcium, sodium, iron, zinc and copper) (Jayanty et al., 2019; Kowalczewski et al., 2019). Further, numerous other natural substances can also be found in potato tubers, including lipids, organic acids, polyphenols, fiber, and glycoalkaloids (Kowalczewski et al., 2019). All of these compounds are also present in potato tuber juice (Fang et al., 2011; Kowalczewski et al., 2012). Thanks to its high nutritional value, potato tuber juice has already been considered a potential substrate for the *Lactobacillus casei* biomass production, as a food additive, or as an ingredient for functional food production (Han et al., 2004; Kim et al., 2012). The biological activity of PJ can also be of value; it can be used in the treatment of bowel diseases, reducing inflammatory symptoms (Kowalczewski et al., 2019).

In commercial media dedicated for BC production, such as HS, usually one specific sugar, typically glucose, is supplied as the carbon source (Jozala et al., 2015; Sperotto et al., 2021). However, in the case of media prepared from waste substrates, a more complex composition and the presence of different sugars should be expected (Hussain et al., 2019). Our analysis of sugar content in PJ medium showed the presence of sucrose (4.0 g/L), glucose (3.5 g/L), fructose (2.4 g/L) and starch (<0.1 g/L) in PJ diluted with water in 1:1 ratio. Thus, in comparison to HS medium which has 20 g/L of glucose, the concentration of glucose alone and the total sugar content in the PJ medium was clearly lower (3.5 g/L and 9.9 g/L, respectively). However, our analysis indicated that diluted PJ contained 25% higher concentration of proteins than HS medium (1.62 g/L v. 1.33 g/L). Finally, the initial pH of PJ medium (6.08) was comparable to that of HS medium (6.19) - neither required any further adjustment of this parameter.

### 3.2. Quantification of K. xylinus cells and BC yield

The kinetics of bacterial growth and the dynamics of BC production constitute crucial factors for adjusting the optimal time and conditions of bacterial cultivation. As a result, both have a major impact on the economic aspect of the bioprocess and its scale up (Abdelraof et al., 2019). In **Fig. 2A** the growth curves of *K. xylinus* in HS and PJ media are presented. As can be seen, the growth curves are comparable. Importantly, for both media, the *K. xylinus* began to grow without a distinct lag phase, indicating that this microorganism did not require a phase adjustment for replication. Molina-Ramirez et al. (2017) reported that the growth curve of cellulose-synthesizing bacteria correlates with affinity for the carbon source. Therefore, here the affinity of *K. xylinus* ATCC 53524 for the carbon sources present in PJ medium was similar to glucose, the only carbon source in HS medium.

**Fig. 2.**
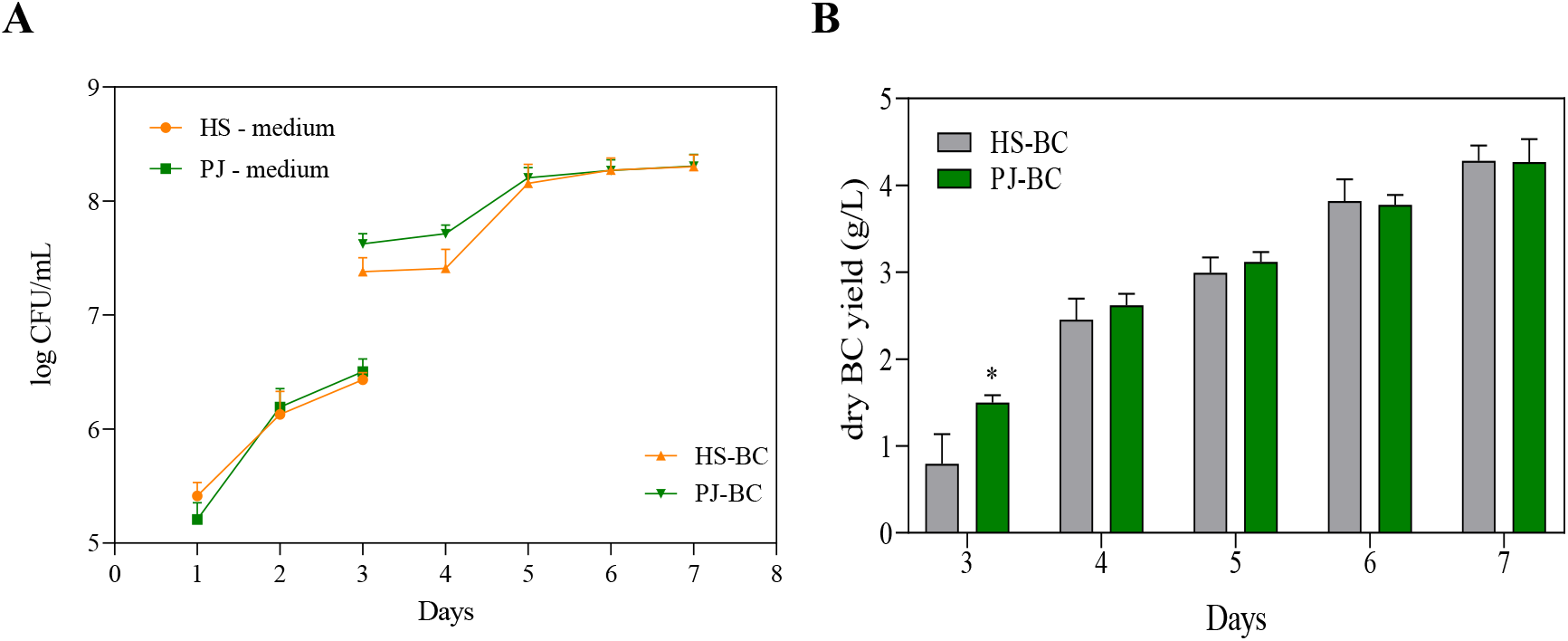
**A)** Number of *K. xylinus* ATCC 53524 cells in HS and PJ media and in BC pellicles. **B)** Yield of BC synthesized by *K. xylinus* ATCC 53524 in HS and PJ media. Data are presented as a mean ± standard error of the mean (SEM). HS - Hestrin-Schramm medium, PJ - potato juice medium, HS-BC - cellulose synthesized in HS, PJ-BC - cellulose synthesized in PJ.

The most important result of the present study was the yield of BC, which - despite the substantial differences in chemical composition of PJ and HS media - was comparable (4.3 g/L) (**Fig. 2B**). It can be noted that on the 3^rd^ day of cultivation in PJ medium, the BC yield was significantly higher. This is likely due to a corresponding higher bacterial cell density (**Fig. 2A**).

As already mentioned, we supplemented both our developed PJ medium and the control HS medium with 1% (v/v) ethanol. The effect of ethanol on BC yield was comparable regardless of the type of medium (**Supplementary material**). Further, we show that for the most efficient production of the BC, potato juice should be diluted with water (**Supplementary material**). Such a result is of particular importance considering the industrial starch production process, in which, depending on the technology used, potato juice can be diluted with water during starch separation or purification (Fang et al., 2011).

### 3.3. The changes in pH and chemical composition of PJ during fermentation process

It is well established that *K. xylinus* can metabolize a variety of sugars, regardless of the presence of the preferred one (Singhsa et al., 2018). Glucose is one of the main components of the HS medium and is the primary substrate for the synthesis of cellulose (**Fig. 3A**). It is incorporated in a 4-step metabolic pathway. First, in the cytosol, glucose is converted to UDP-glucose by uridine-diphosphate-glucose pyrophosphorylase (EC 2.7.7.64, UDP-glucose pyrophosphorylase). In the next step, cellulose synthase (EC 2.4.1.12), located in the bacterial cell wall, catalyzes the polymerization of UDP-glucose to poly β-1-4 glucan (Lei et al., 2012). However, although the BC is an anhydrous polymer of glucose, *K. xylinus* can use other monosaccharides and disaccharides, such as fructose, xylitol, sucrose, maltose or lactose (Chen et al., 2019). This feature is also apparent in the present study (**Fig. 3B**). Importantly, the use of glucose as the primary carbon source for *K. xylinus* is associated with a drawback. These bacteria can readily convert glucose to gluconic acid, significantly decreasing the pH (< 3.5) of the medium (**Fig. 3C**). This, in turn, results in a decrease of intracellular pH. As a result, high concentration of gluconic acid can change the activity of metabolic pathways responsible for BC synthesis, reducing the yield (Aswini et al., 2020; Embuscado et al., 1994). In contrast, when the culture medium contains fructose, *K. xylinus* cells convert this monosaccharide to acetic acid, which results a lower drop in pH, as compared to gluconic acid (Drozd et al., 2021). Therefore, the use of PJ, containing a mixture of sugars, including fructose, with a relatively low concentration of glucose (3.5 g/L) as compared to the HS (20 g/L), resulted in a significantly reduced concentration of gluconic acid produced during fermentation and the pH remained above 4, even at the end to the fermentation process (**Fig. 3D**). Meanwhile, in cultures with HS medium, the concentration of gluconic acid at the end of the fermentation process was approx. 10 times higher and the pH dropped to 3.5 (**Fig. 3C,D**).

**Fig. 3.**
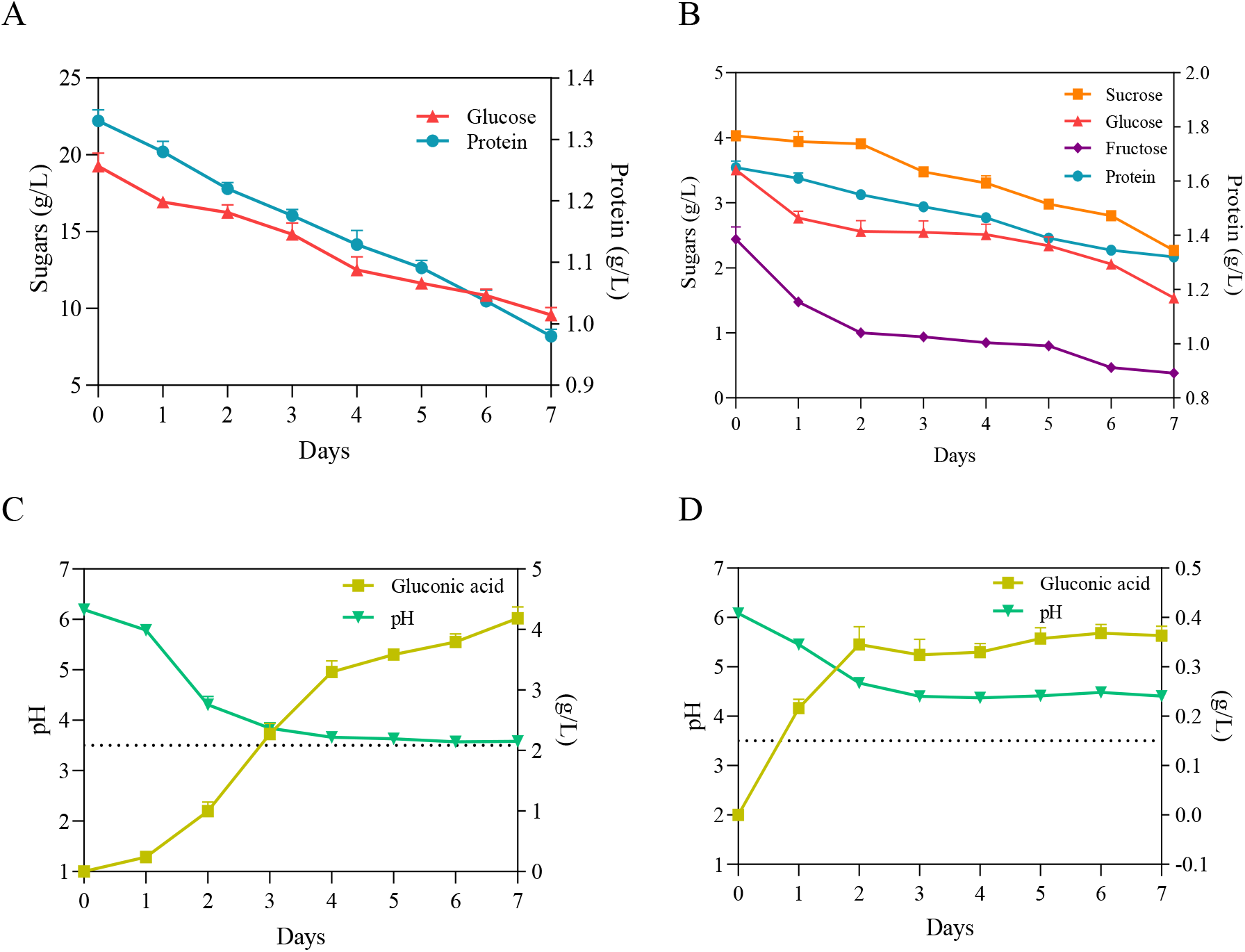
**A)** Concentration of glucose and protein in HS medium. **B)** Concentration of sugars and protein in PJ medium. **C)** pH and gluconic acid concentration in HS medium. **D)** pH and gluconic acid concentration in PJ medium - all before and during BC biosynthesis by *K. xylinus* ATCC 53524. Data are presented as a mean ± standard error of the mean (SEM). ^…^ critical pH value (3.5).

Nitrogen is a key component of proteins, making it necessary for cell metabolism. It comprises 8-14% of dry cell mass of bacteria. Alongside carbon sources, proteinaceous nitrogen sources can also contribute to the enhancement of biomass production and BC synthesis, if suitably chosen (Ramana and Singh, 2000). The reports by several authors have shown that *K. xylinus* strains can utilize a wide range of protein and nitrogen sources, including casein hydrolysate, peptone, corn steep liquor, yeast extract, glutamate, soybean meal, glycine, and ammonium sulphate (Chawla et al., 2009; Ramana and Singh, 2000). Likewise, in our present study, we observed that, although PJ and HS media differed in terms of qualitative and quantitative protein content, protein consumption during the BC biosynthesis process was similar (0.33 g/L v. 0.35 g/L) (**Fig. 3A,B**).

The present study also confirmed that the starch was not metabolized during BC biosynthesis process, as its concentration, 0.67 g/L, in media did not change during cultivation of *K. xylinus*. Further, when using non-centrifuged PJ, containing starch solids, BC yield significantly decreased (**Supplementary material**).

### 3.4. The role of bacterial strain and presence of sugars on BC yield

The process of BC synthesis by bacterial cells is multilevel and includes many, tightly connected, metabolic pathways. A recent phylogenetic study of *Komagataeibacter* strains with defined genomes suggests that there is high variability in the structure of the pathways involved in BC synthesis (Drozd et al., 2021). The differences in structure and number of cellulose synthase operons in the genomes of *Komagataeibacter* strains are also considered the main reason behind inter-species differences in BC productivity and response to cultivation conditions (La China et al., 2020). Therefore, ideally the culture medium used to produce BC should be adapted to the specific bacterial strain prior the fermentation process (Drozd et al., 2021; La China et al., 2020). On the other hand, considering the results reported by other authors related to the optimization of the culture medium, it can be noted that culture media with a more complex composition, particularly in terms of carbon and nitrogen sources, are more universal and result in relatively high yields of BC regardless of the bacterial strain used (Chawla et al., 2009; Molina-Ramírez et al., 2017). For this reason, we also aimed to determine whether PJ medium can be considered universal for the cultivation of different *K. xylinus* strains. In 4 out of the 8 tested strains (**Fig. 4A**), BC yield using PJ medium was equivalent to HS medium.

**Fig. 4.**
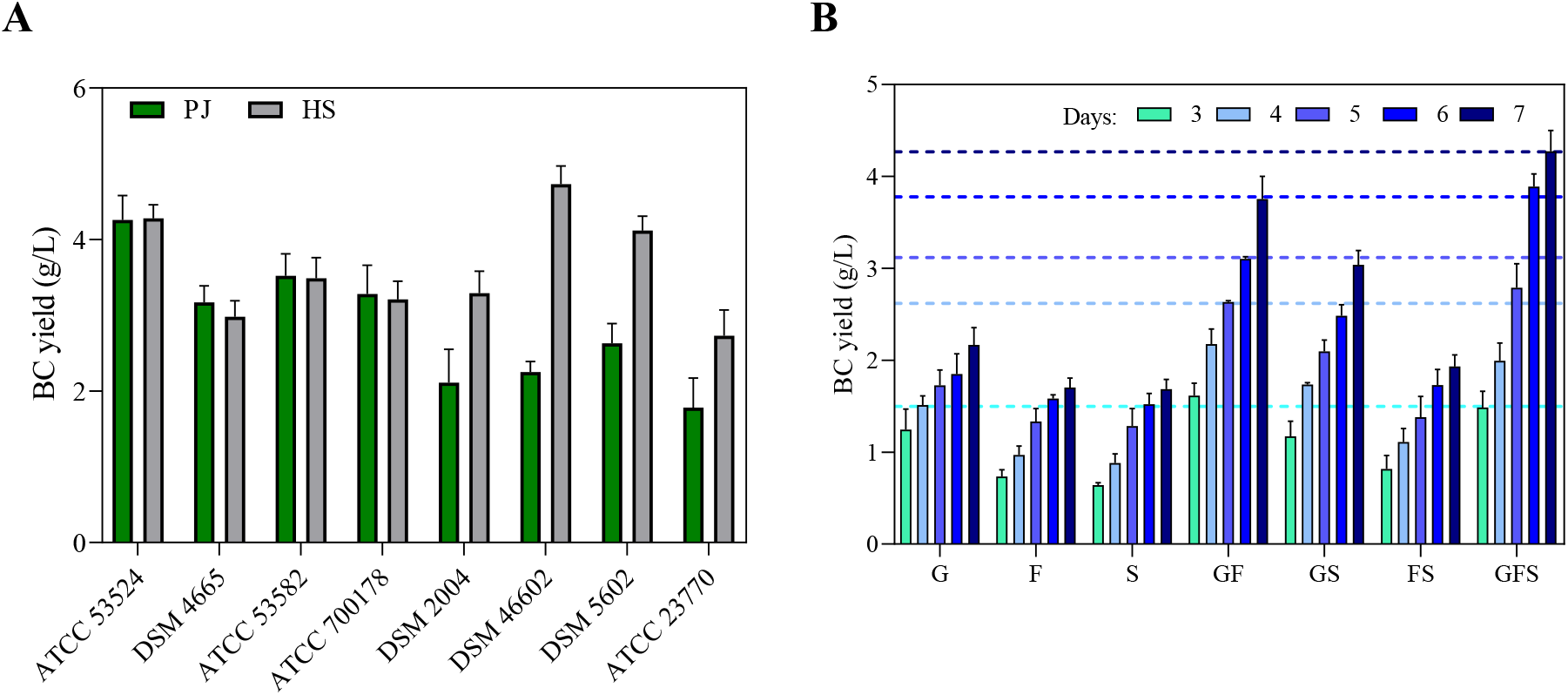
**A)** BC yields of different *K. xylinus* strains in PJ and HS media after 7 days of cultivation. **B)** BC yields of *K. xylinus* ATCC 53524 in HS medium with the addition of sugars at the same concentration as in PJ medium in the following days of cultivation. Data are presented as a mean ± standard error of the mean (SEM). G - glucose, F - fructose, S - sucrose. Horizontal lines in (B) indicate BC yield of *K. xylinus* ATCC 53524 in PJ medium in the following days of cultivation.

Further, in order to determine which of the carbon sources present in the PJ plays a crucial role for the high BC yield, we prepared HS media supplemented with glucose, fructose, sucrose or a combination of these sugars at the same concentrations as in PJ. The results showed that all of the sugars present in PJ medium were metabolized and converted into BC by the *K. xylinus* ATCC 53524 strain (**Fig. 4B**). The lowest BC yield was obtained when only a single sugar was present in the medium. Higher BC yields were observed for glucose-fructose and glucose-sucrose combinations. However, the highest BC yield (4.27 g/L) was obtained for the combination of glucose, fructose, and sucrose, that reflect the composition of the main sugars present in the PJ (**Fig. 3B v. Fig. 4B**).

### 3.5. Physicochemical properties of BC obtained from potato tuber juice medium

#### 3.5.1. Macro- and microstructure

Macromorphological assessment of PJ-BC indicated that it has a homogenous, smooth surface with no visible residues of the culture medium. In fact, the micromorphology was similar to the typical micromorphology of mature BC synthesized in HS medium (**Fig. 5A,B,D,E**). Likewise, the microstructure of PJ-BC was similar to that of HS-BC (**Fig. 5C,F**) and was consistent with that described in the literature (Klemm et al., 2001). These observations are particularly important given that the PJ medium used is typically a waste product. A frequent issue encountered with the use of other agroindustrial wastes such as fruit and vegetable peels or juices (besides the previously discussed issues of pre-processing and/or insufficient nutritional content), is the high content of artefacts, solids, and/or pigments present in the raw substrates. These will then also be present in the medium and then also in the BC obtained.

**Fig. 5.**
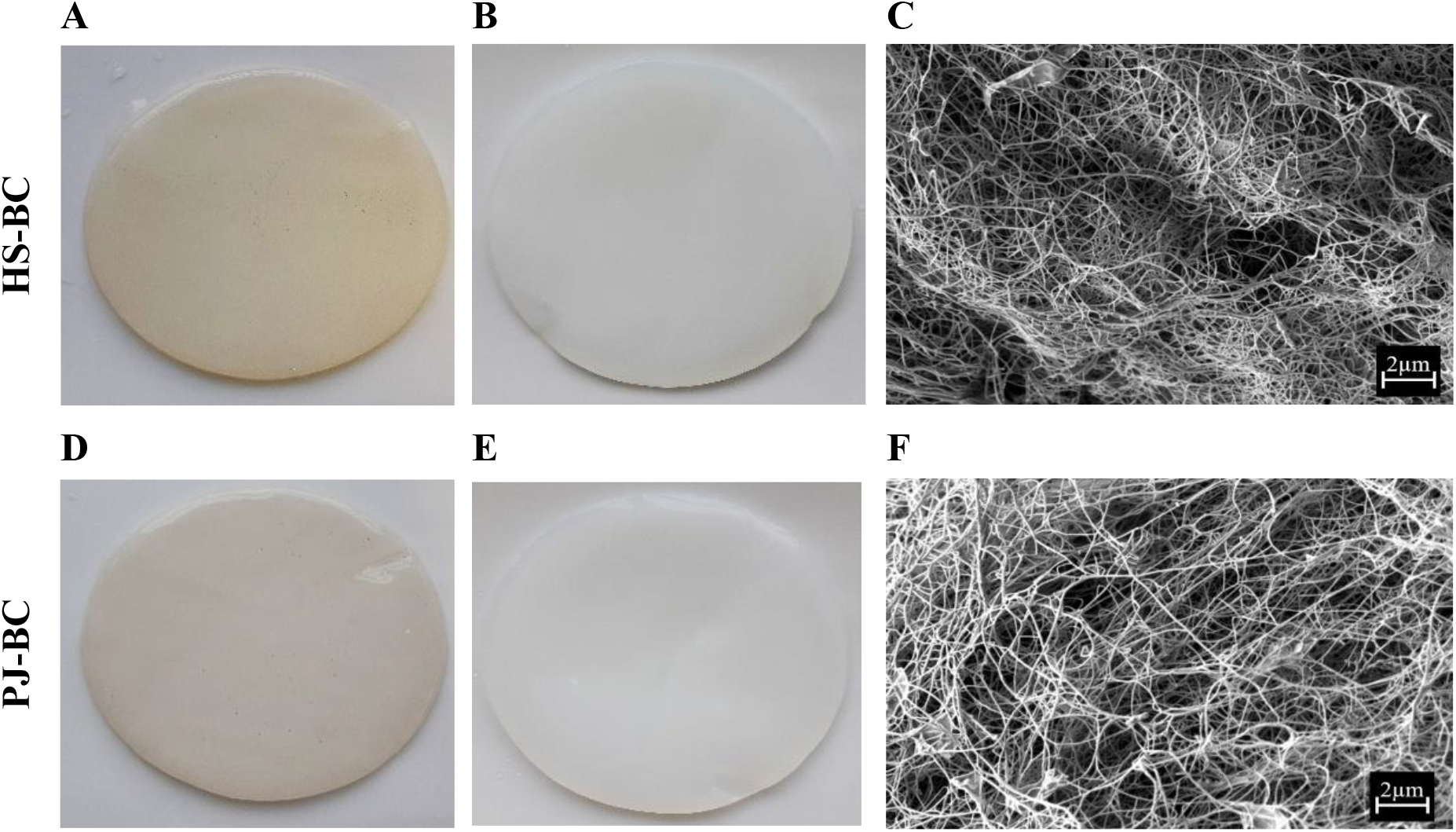
Macromorphology of BC from *K. xylinus* ATCC 53524: **A, D** - before purification, **B, E** - after purification. **C, F** - micromorphology of BC from *K. xylinus* ATCC 53524 after purification, magnification 10000x (SEM, Auriga 60, Zeiss, Oberkochen, Germany).

In order to assess the potential advantage of PJ over the other natural ingredient-based media, we prepared media from other food/agro-industry waste (potato peels, orange peels, beetroots, and apples) in the same fashion as the PJ media. We noted that, in terms of transparency, PJ medium was comparable to HS medium. Further, for the case of PJ, there was less solids that had to be removed prior the BC production. Representative pictures of the tested media and BC membranes obtained are presented in the **Supplementary material**. As was anticipated, the colors of the media prepared from the beetroots, apples, and even potato peels were also reflected in the pigmentation of the BC pellicles. This necessitated a longer purification process, as compared to BC obtained from cultures in PJ or HS media. In the case of BC obtained from the orange peel-based medium, the yellow pigment was removed in the first stage of purification. However, this relative ease of pigment removal likely due to the reduced thickness of the obtained cellulose membrane and its amorphous structure, which reduce its potential industrial applicability. In the case of other fruit-based media, some residues of fruits were tightly bound to the BC and remained, even after the purification process, see the **Supplementary material**. In contest, none of these issues were observed when PJ medium was used for BC synthesis.

#### 3.5.2. Analysis of ATR-FTIR spectra

To further compare the BC obtained from PJ medium cultures to those in HS, we used infrared spectroscopy to examine chemical composition and crystallinity. The ATR-FTIR spectra of both HS-BC and PJ-BC exhibited all of the characteristic adsorption bands of cellulose functional groups (**Supplementary material**) that have been previously reported by other authors (Algar et. al., 2015; Huang et al., 2010). In all of the spectra, a pair of closely located bands at 710 cm^-1^ and 750 cm^-1^ was clearly visible and can be assigned to *Iα* and *Iβ* of BC microfibril allomorphs. All of the BC samples, regardless the medium used, displayed a relatively high content of Iα fraction (**Tab. 1**).

**Tab. 1.** Selected parameters of BC pellicles synthesized by *K. xylinus* ATCC 53524 in HS and PJ media.

|  | **SR** | **WHC** | **EB** | **TS** |
| --- | --- | --- | --- | --- |
| **HS-BC** | 299 ± 33 | 3.70 ± 1.24 | 20.0 ± 3.67 | 2.00 ± 0.26 |
| **PJ-BC** | 298 ± 33 | 4.07 ± 1.45 | 19.80 ± 3.28 | 2.05 ± 0.23 |
|  | **Iα fraction** | **I.C.1<sub>1370/2900</sub>** | **I.C.2<sub>1430/900</sub>** | **$\rho$** |
| **HS-BC** | 0.44 ± 0.013 | 1.62 ± 0.21 | 1.04 ± 0.06 | 1.55 ± 0.02 |
| **PJ-BC** | 0.45 ± 0.018 | 1.57 ± 0.27 | 1.10 ± 0.11 | 1.49 ± 0.07 |

Data are presented as a mean ± standard error of the mean (SEM). SR – swelling ratio (%); WHC – water holding capacity after 8 min at 60°C (%); EB - Elongation at break (%); TS - Tensile strength (MPa); $\rho$ - density of BC ( $\text{g/m}^3$ ).

Importantly, the use of PJ medium did not influence the crystallinity indexes of the obtained BC, as compared to using HS. All of the samples exhibited high values of I.C. I1430/900. Likewise, the second crystallinity index, calculated as a ratio between the absorbance of the bands at 1370 cm^-1^ (CH bending) and 2900 cm^-1^ (CH and CH2 stretching) was also similar, regardless the medium used.

#### 3.5.3. Water-related properties, density, and mechanical properties of BC

For many applications, the water-related properties can be considered to be the most important feature of BC, because they determine its absorption capacity and ability to retain and release liquid. Overall, results (**Tab. 1, Supplementary material**) showed that water-related parameters were comparable for BC membranes regardless the medium used for their production. There were no statistically significant differences between results of the total swelling ratio and water holding capacity. Likewise, regardless of culture medium, we observed no significant differences in density nor tensile mechanical properties (**Tab. 1, Supplementary material**). All of the obtained results were in good agreement with values reported in the literature for typical, unmodified BC (Markiewicz et al., 2004).

### 3.6. BC cytotoxicity screening

BC has numerous potential biomedical applications including use as a biomaterial in tissue engineering, wound healing, and drug delivery (Czaja et al., 2006; Sannino et al., 2009). In this context, it’s important to note that BC is considered non-toxic. Therefore, it was important to confirm that BC obtained from PJ medium cultures was also not cytotoxic. We conducted extract and direct contact *in vitro* cytotoxicity assays, based on ISO10993-5. In the extract tests, we observed robust growth of L929 cells exposed to all extracts. None of the tested materials resulted in viability below 70%, the threshold for cytotoxicity according to ISO10993-5 (**Fig. 6A**). There was no difference between PJ-BC and HS-BC. A similar trend was also observed in the direct contact assay. Importantly, the results of the viability assay were directly confirmed using fluorescence microscopy, with only live (stained green) cells observed (**Fig. 6B**). Likewise, the morphology of L929 cells was not altered by either BC (**Supplementary material**). The results are in good agreement with our previous studies on BC based materials (Junka et al., 2017). However, we cannot compare our results to BC obtained other by-product or waste media (Hussain et al., 2019; Kongruang, 2007; Revin et al., 2018), because cytotoxicity studies are not (yet) a standard research practice in this context. We conclude that, from a cytotoxicity standpoint, BC obtained from PJ medium is equivalent to that obtained from conventional culture, making it suitable for biomedical applications.

**Fig. 6.**
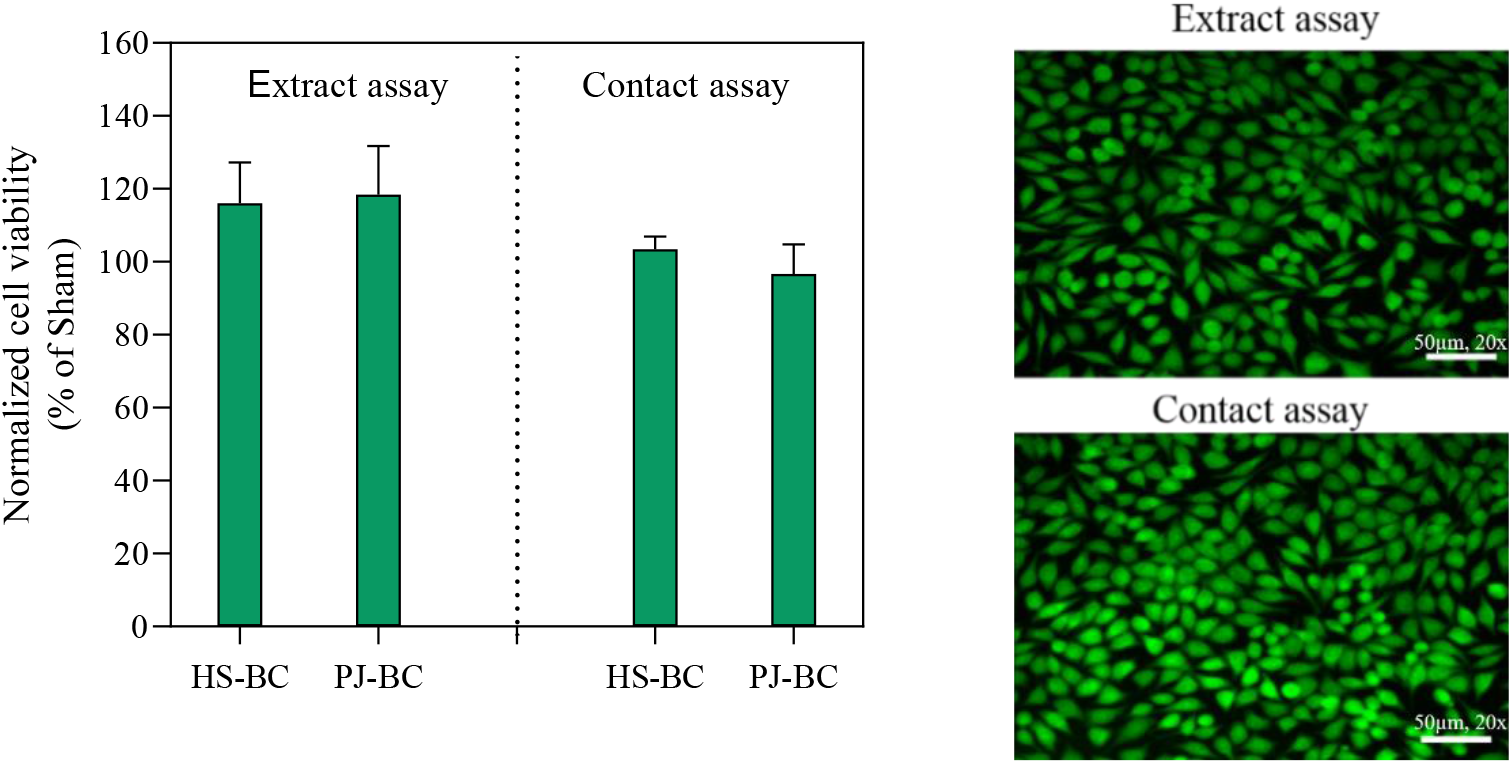
**A)** L929 fibroblast viability data, as percentage of sham. **B)** Fluorescence Live/Dead visualization of L929 fibroblasts exposed PJ-BC for 24 hours (extract and direct contact) and stained with Syto 9 (green, live) and propidium iodide dyes (red, dead). Data are presented as mean ± standard error of the mean (SEM); HS-BC - BC biosynthesized in HS medium, PJ-BC - BC biosynthesized in PJ medium.

## SUMMARY

Our results demonstrate that the PJ (without any pre-treatment) is suitable as source of nutrients for cellulose-producing *K. xylinus* bacteria. Most importantly, after diluting PJ with water 1:1 to prepare medium, the yield of BC was equivalent to that obtained from commercial, HS medium. Further, the PJ-BC obtained in this study did not differ from conventionally produced HS-BC in terms of its physical and chemical properties and was not cytotoxic. As a result, PJ-BC should be able to be used in the same applications as commercially produced BC. Importantly, converting the BC production process to use PJ medium at an industrial scale should be relatively easy to implement, thanks to the ready, high availability of PJ, a by-product of the potato starch industry.

## References

1. Abdelraof, M., Hasanin, M.S., El-Saied, H., 2019. Ecofriendly green conversion of potato peel wastes to high productivity bacterial cellulose. Carbohydr. Polym. 211, 75–83.

2. Abol-Fotouh, D., Hassan, M.A., Shokry, H., Roig, A., Azab, M.S., Kashyout, A.B., 2020. Bacterial nanocellulose from agro-industrial wastes: low-cost and enhanced production by *Komagataeibacter saccharivorans* MD1. Sci. Rep. 10(1), 1–14.

3. Algar, I., Fernandes, S.C., Mondragon, G., Castro, C., Garcia-Astrain, C., Gabilondo, N., Retegi, A., Eceiza, A., 2015. Pineapple agroindustrial residues for the production of high value bacterial cellulose with different morphologies. J. Appl. Polym. Sci. 132(1), 1–17.

4. Aswini, K., Gopal, N.O., Uthandi, S., 2020. Optimized culture conditions for bacterial cellulose production by *Acetobacter senegalensis* MA1. BMC Biotechnol. 20(1), 1–16.

5. Bradford, M.M., 1976. A rapid and sensitive method for the quantitation of microgram quantities of protein utilizing the principle of protein-dye binding. Anal. Biochem. 72(1-2), 248–254.

6. Bradshaw, J.E., Bonierbale, M., 2010. Potatoes, in: Bradshaw, J.E. (Ed.), Root and tuber crops. Springer, New York, pp. 1–52.

7. Chawla, P.R., Bajaj, I.B., Survase, S.A., Singhal, R.S., 2009. Microbial cellulose: fermentative production and applications. Food Technol. Biotechnol. 47(2), 107–124.

8. Chen, G., Wu, G., Chen, L., Wang, W., Hong, F.F., Jönsson, L.J., 2019. Comparison of productivity and quality of bacterial nanocellulose synthesized using culture media based on seven sugars from biomass. Microb. Biotechnol. 12(4), 677–687.

9. Cheng, Z., Yang, R., Liu, X., Liu, X., Chen, H., 2017. Green synthesis of bacterial cellulose via acetic acid pre-hydrolysis liquor of agricultural corn stalk used as carbon source. Bioresour. Technol. 234, 8–14.

10. Ciecholewska-Juśko, D., Żywicka, A., Junka, A., Drozd, R., Sobolewski, P., Migdał, P., Kowalska, U., Toporkiewicz, M., Fijałkowski, K., 2021. Superabsorbent crosslinked bacterial cellulose biomaterials for chronic wound dressings. Carbohydr. Polym. 253, 1–13.

11. Czaja, W., Krystynowicz, A., Bielecki, S., Brown Jr, R.M., 2006. Microbial cellulose - the natural power to heal wounds. Biomaterials. 27(2), 145–151.

12. Drozd, R., Szymańska, M., Żywicka, A., Kowalska, U., Rakoczy, R., Kordas, M., Konopacki, M., Junka, A.F., Fijałkowski, K., 2021. Exposure to non-continuous rotating magnetic field induces metabolic strain-specific response of *Komagataeibacter xylinus*. Biochem. Eng. J. 166, 1–12.

13. Embuscado, M.E., Marks, J.S., BeMiller, J.N., 1994. Bacterial cellulose. I. Factors affecting the production of cellulose by Acetobacter xylinum. Food Hydrocoll. 8(5), 407–418.

14. Fang, C., Boe, K., Angelidaki, I., 2011. Biogas production from potato-juice, a by-product from potato-starch processing, in upflow anaerobic sludge blanket (UASB) and expanded granular sludge bed (EGSB) reactors. Bioresour. Technol. 102(10), 5734–5741.

15. Goelzer, F.D.E., Faria-Tischer, P.C.S., Vitorino, J.C., Sierakowski, M.R, Tischer, C.A., 2009. Production and characterization of nanospheres of bacterial cellulose from *Acetobacter xylinum* from processed rice bark. Mat. Sci. Eng. C. 29(2), 546–551.

16. Grommers, H.E., van der Krogt, D.A., 2009. Potato starch: production, modifications and uses. Starch. 11, 511–539.

17. Han, G.P., Lee, K.R., Han, J.S., Kozukue, N., Kim, D.S., Kim, J., Bae, J.H., 2004. Quality characteristics of the potato juice-added functional white bread. Korean J. Food Sci. Technol. 36(6), 924–929.

18. Huang, H.C., Chen, L.C., Lin, S.B., Hsu, C.P., Chen, H.H., 2010. *In situ* modification of bacterial cellulose network structure by adding interfering substances during fermentation. Bioresour. Technol. 101(15), 6084–6091.

19. Hussain, Z., Sajjad, W., Khan, T., Wahid, F., 2019. Production of bacterial cellulose from industrial wastes: a review. Cellulose 26(5), 2895–2911.

20. Jayanty, S.S., Diganta, K., Raven, B., 2019. Effects of cooking methods on nutritional content in potato tubers. Am. J. Potato Res. 96(2), 183–194.

21. Jozala, A.F., Pértile, R.A.N., dos Santos, C.A., de Carvalho Santos-Ebinuma, V., Seckler, M.M., Gama, F.M., Pessoa, A., 2015. Bacterial cellulose production by *Gluconacetobacter xylinus* by employing alternative culture media. Appl. Microbiol. Biotechnol. 99(3), 1181–1190.

22. Junka, A., Fijałkowski, K., Ząbek, A., Mikołajewicz, K., Chodaczek, G., Szymczyk, P., Smutnicka, D., Żywicka, A., Sedghizadeh, P.P., Dziadas, M., Młynarz, P., Bartoszewicz, M., 2017. Correlation between type of alkali rinsing, cytotoxicity of bio-nanocellulose and presence of metabolites within cellulose membranes. Carbohydr. Polym. 157, 371–379.

23. Kataoka, Y., Kondo, T., 1999. Quantitative analysis for the cellulose *Iα* crystalline phase in developing wood cell walls. Int. J. Biol. Macromol. 24(1), 37–41.

24. Kim, N.J., Jang, H.L., Yoon, K.Y., 2012. Potato juice fermented with *Lactobacillus casei* as a probiotic functional beverage. Food Sci. Biotechnol. 21(5), 1301–1307.

25. Klemm, D., Schumann, D., Udhardt, U., Marsch, S., 2001. Bacterial synthesized cellulose artificial blood vessels for microsurgery. Prog. Polym. Sci. 26(9), 1561–1603.

26. Kongruang, S., 2007. Bacterial cellulose production by *Acetobacter xylinum* strains from agricultural waste products. Biotechnol. Fuels Chem. 763–774.

27. Kurosumi, A., Sasaki, C., Yamashita, Y., Nakamura, Y., 2009. Utilization of various fruit juices as carbon source for production of bacterial cellulose by *Acetobacter xylinum* NBRC 13693. Carbohydr. Polym. 76(2), 333–335.

28. Kot, A.M., Pobiega, K., Piwowarek, K., Kieliszek, M., Błażejak, S., Gniewosz, M., Lipińska, E., 2020. Biotechnological methods of management and utilization of potato industry waste - a review. Potato Res. 63, 431–447.

29. Kowalczewski, P., Celka, K., Białas, W., Lewandowicz, G., 2012. Antioxidant activity of potato juice. In Acta Sci. Pol. Technol. Aliment. 11(2), 175–181.

30. Kowalczewski, P.Ł., Olejnik, A., Białas, W., Rybicka, I., Zielińska-Dawidziak, M., Siger, A., Kubiak P., Lewandowicz, G., 2019. The nutritional value and biological activity of concentrated protein fraction of potato juice. Nutrients 11(7), 1523–1537.

31. La China, S., Bezzecchi, A., Moya, F., Petroni, G., Di Gregorio, S., Gullo, M., 2020. Genome sequencing and phylogenetic analysis of K1G4: a new *Komagataeibacter* strain producing bacterial cellulose from different carbon sources. Biotechnol. Lett. 42(5), 807–818.

32. Lei, L., Li, S., Gu, Y., 2012. Cellulose synthase complexes: composition and regulation. Front. Plant Sci. 3, 75–84.

33. Li, Z., Wang, L., Hua, J., Jia, S., Zhang, J., Liu, H., 2015. Production of nano bacterial cellulose from waste water of candied jujube-processing industry using *Acetobacter xylinum*. Carbohydr. Polym. 120, 115–119.

34. Lima, H.L.S., Nascimento, E.S., Andrade, F.K., Brígida, A.I.S., Borges, M. de F., Cassales, A.R., Muniz, C.R., Souza Filho, M. De S.M., Morais, J.P.S., Rosa, M. de F., 2017. Bacterial cellulose production by *Komagataeibacter hansenii* ATCC 23769 using sisal juice - an agroindustry waste. Braz. J. Chem. Eng. 34(3), 671–680.

35. Markiewicz, E., Hilczer, B., Pawlaczyk, C., 2004. Dielectric and acoustic response of biocellulose. Ferroelectrics. 304(1), 39–42.

36. Molina-Ramírez, C., Castro, M., Osorio, M., Torres-Taborda, M., Gómez, B., Zuluaga, R., Gómez, C., Ganan, P., Rojas, O.J., Castro, C., 2017. Effect of different carbon sources on bacterial nanocellulose production and structure using the low pH resistant strain *Komagataeibacter medellinensis*. Materials. 10(6), 639.

37. Ramana, K.V., Tomar, A., Singh, L., 2000. Effect of various carbon and nitrogen sources on cellulose synthesis by *Acetobacter xylinum*. World J. Microbiol. Biotechnol. 16(3), 245–248.

38. Revin, V., Liyaskina, E., Nazarkina, M., Bogatyreva, A., Shchankin, M., 2018. Cost-effective production of bacterial cellulose using acidic food industry by-products. Braz. J. Med. Biol. Res. 49, 151–159.

39. Riss, T.L., Moravec, R.A., Niles, A.L., Duellman, S. Benink, H.A., Worzella, T.J., Minor, L., 2004. Cell Viability Assays, in: Sittampalam, G., Coussens, N. (Eds.), Assay Guidance Manual. Eli Lilly & Company and the National Center for Advancing Translational Sciences, Bethesda, MD.

40. Sannino, A., Demitri, C., Madaghiele, M., 2009. Biodegradable cellulose-based hydrogels: design and applications. Materials. 2(2), 353–373.

41. Singhsa, P., Narain, R., Manuspiya, H., 2018. Physical structure variations of bacterial cellulose produced by different *Komagataeibacter xylinus* strains and carbon sources in static and agitated conditions. Cellulose 25(3), 1571–1581.

42. Sperotto, G., Stasiak, L.G., Godoi, J.P.M.G., Gabiatti, N.C., De Souza, S.S., 2021. A review of culture media for bacterial cellulose production: complex, chemically defined and minimal media modulations. Cellulose 28, 2649–2673.

43. Sulistyarti, H., Fardiyah, Q., Febriyanti, S., 2015. A simple and safe spectrophotometric method for iodide determination. Makara J. Sci. 19(2), 43–48.

44. Ullah, H., Badshah, M., Correia, A., Wahid, F., Santos, H.A., Khan, T., 2019. Functionalized bacterial cellulose microparticles for drug delivery in biomedical applications. Curr. Pharm. Des. 25(34), 3692–3701.

45. https://starch.eu/the-european-starch-industry/ (May, 2021)

