## Supplementary material for "Potato juice, a starch industry waste, as a cost-effective medium for the biosynthesis of bacterial cellulose"

**Corresponding author**

**Tab. S1.** Dry BC yield of *K. xylinus* ATCC 53524 depending on the degree of potato juice dilution with water, addition of ethanol and starch content.

| | Dry BC yield (g/L) |
|---|---|
| PJ:water dilution ratio 1:1* | 4.26 ± 0.32 |
| PJ:water dilution ratio 1:2 | 1.97 ± 0.11 |
| PJ:water dilution ratio 2:1 | 2.27 ± 0.18 |
| HS with 1% of ethanol | 4.28 ± 0.18 |
| HS without ethanol | 2.04 ± 0.37 |
| PJ with 1% of ethanol* | 4.26 ± 0.32 |
| PJ without ethanol | 1.97 ± 0.53 |
| non-centrifuged PJ medium containing starch solids (with 0.67 g/L starch content) | 1.689 ± 0.143 |
| PJ medium prepared with decantation and centrifugation (with <0.1 g/L starch content)* | 4.26 ± 0.32 |

Data are presented as a mean ± standard error of the mean (SEM). * - these are the same conditions, presented separately for purposes of comparison.

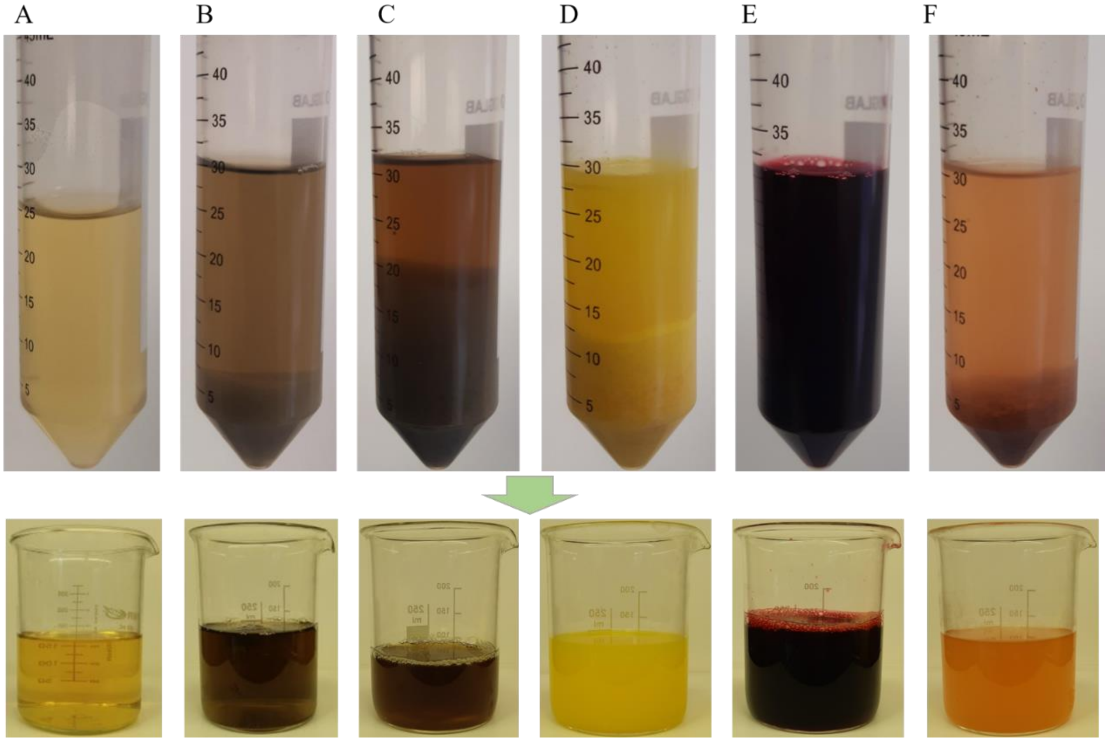


**Fig. S1.** HS and natural ingredients-based media before (in 50 mL plastic tubes) and after decantation (in glass beakers); **A)** HS; **B)** potato juice; **C)** potato peels; **D)** orange peels; **E)** beetroot; **F)** apple.

**A**

| Type of medium | After NaOH treatment | After 24 h of water rinsing | After 48 h of water rinsing |
|---|---|---|---|
| HS | 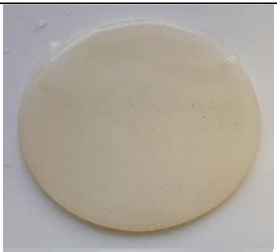 | 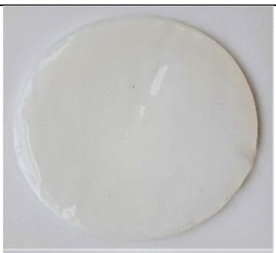 | 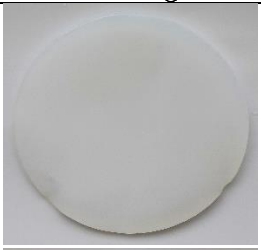 |
| Potato juice | 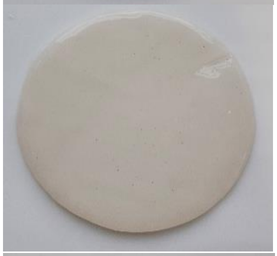 | 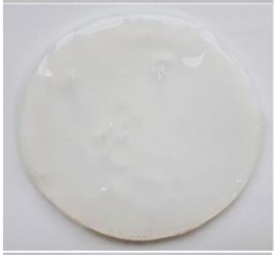 | 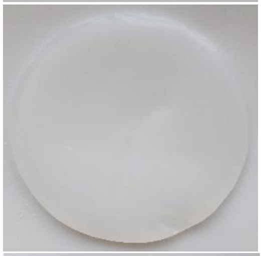 |
| Potato peels | 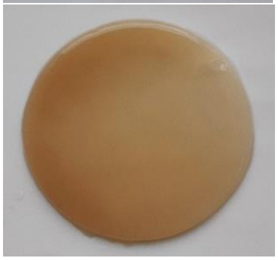 | 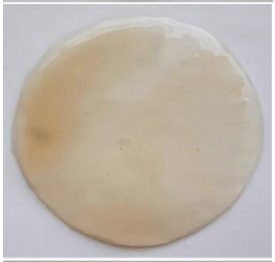 | 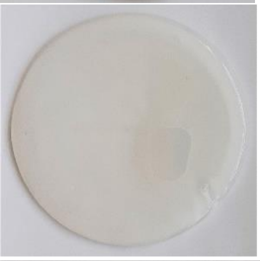 |
| Orange peels | 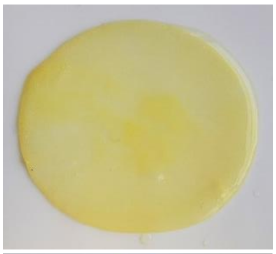 | 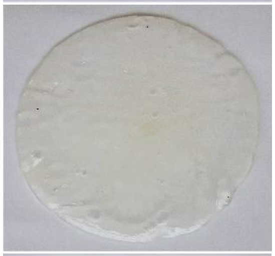 | 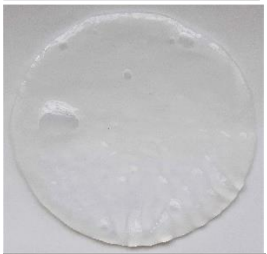 |
| Beetroot | 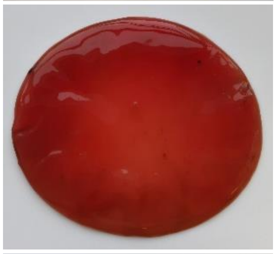 | 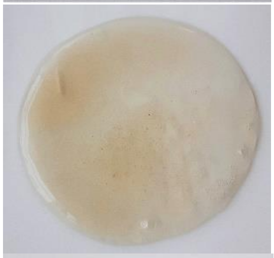 | 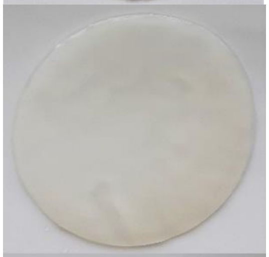 |
| Apple | 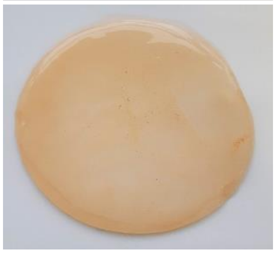 | 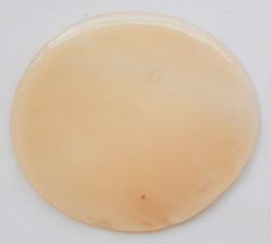 | 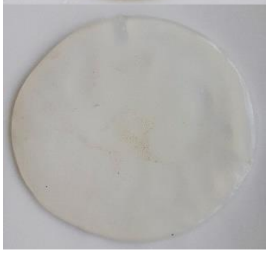 |

**B**

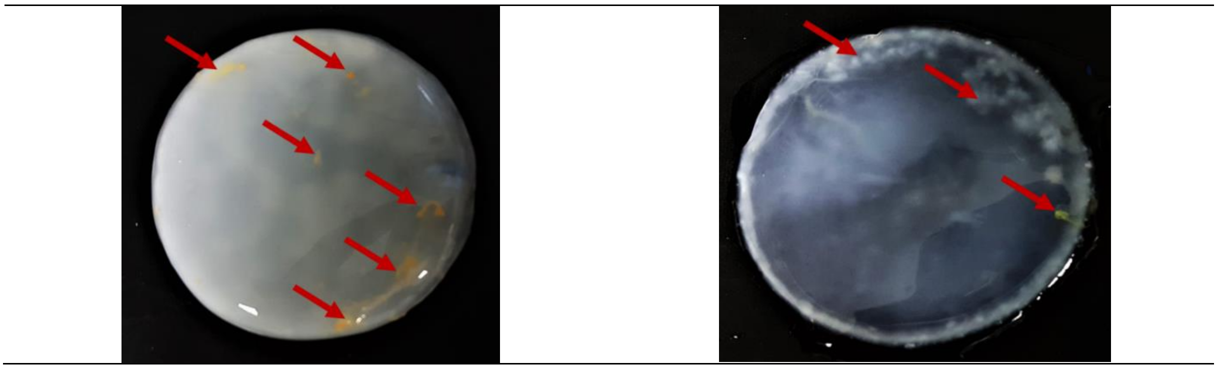

**Fig. S2.** BC obtained from *K. xylinus* ATCC 53524 in HS and natural ingredients-based media in subsequent stages of purification (**A)** Impurities in BC from *K. xylinus* ATCC 53524 synthesized using natural (orange peels) ingredients-based media **(B).**

Red arrows indicate residues fragments (hard-to-remove impurities) of fruits bound in the BC membrane.

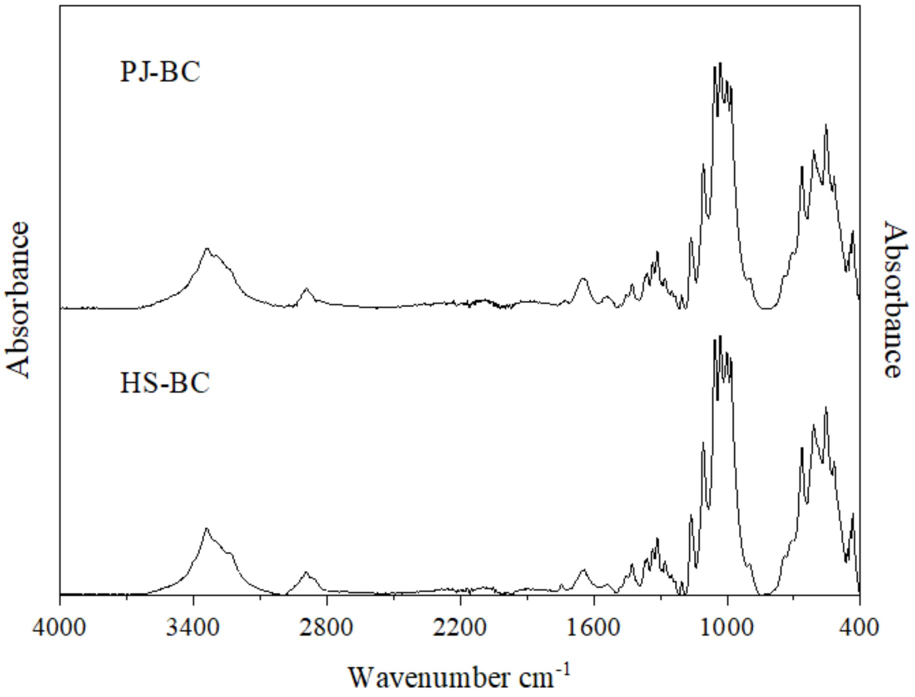


**Fig. S3.** ATR-FTIR spectra of HS-BC and PJ-BC obtained from *K. xylinus* ATCC 53524 cultures.

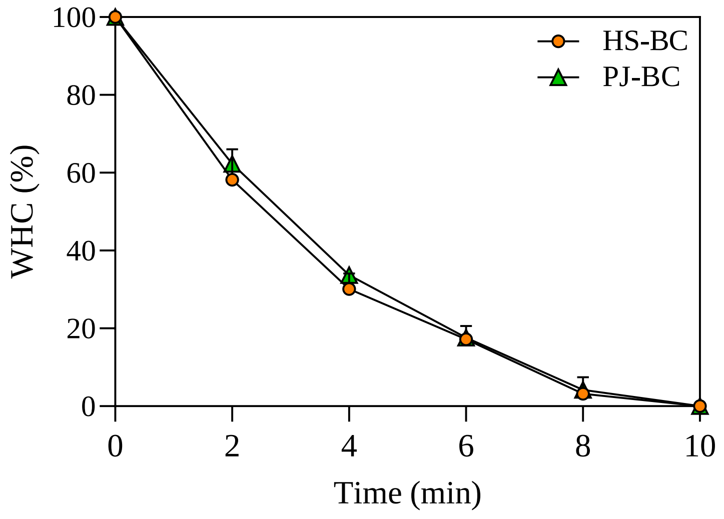


**Fig. S4.** Water holding capacity (%) of HS-BC and PJ-BC obtained from *K. xylinus* ATCC 53524 during drying process at 60°C.

Data are presented as a mean ± standard error of the mean (SEM).

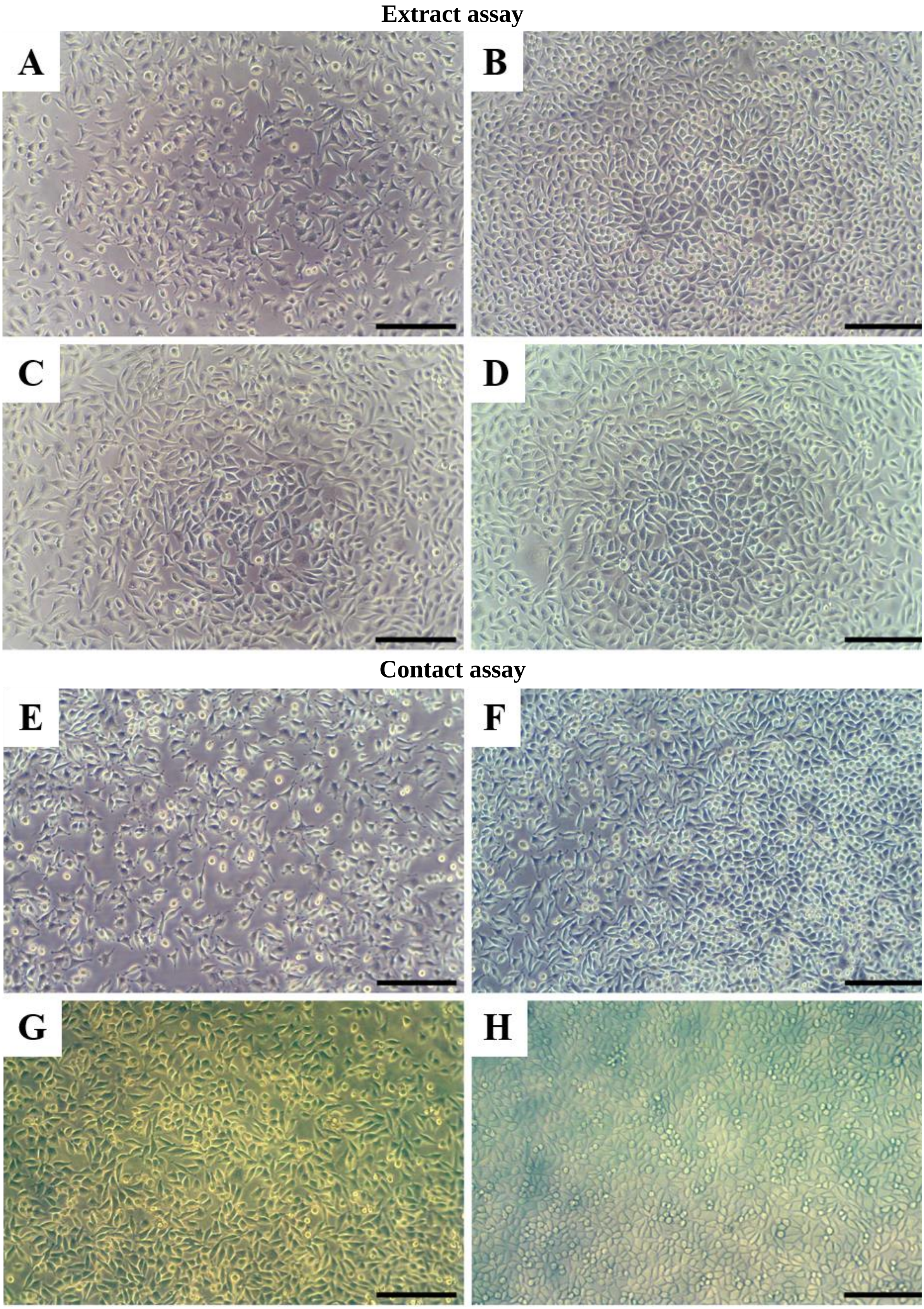


**Fig. S5.** Representative micrographs of L929 cells. **A)** 24 h after seeding (10000 cells per well); **B)** After 24 h of incubation with sham extract; **C, D)** After 24 h incubation with extracts of HS-BC and PJ-BC obtained from *K. xylinus* ATCC 53524, respectively; **E)** 24 h after seeding, prior to disc placement; **F)** After incubation for 24 h without disc (sham); **G, H)** After 24 h incubation beneath discs of HS-BC and PJ-BC obtained from *K. xylinus* ATCC 53524, respectively. Images taken with discs in place - prior to removal. Scale bar indicates 200 µm.
